# Glycolysis–Wnt signaling axis tunes developmental timing of embryo segmentation

**DOI:** 10.1101/2024.01.22.576629

**Authors:** Hidenobu Miyazawa, Jona Rada, Paul Gerald Layague Sanchez, Emilia Esposito, Daria Bunina, Charles Girardot, Judith Zaugg, Alexander Aulehla

**Author notes:** These authors contributed equally to this work (Hidenobu Miyazawa, Jona Rada).

## Abstract

The question of how metabolism impacts development is seeing a renaissance [1, 2]. How metabolism exerts instructive signaling functions is one of the central issues that need to be resolved. We tackled this question in the context of mouse embryonic axis segmentation. Previous studies have shown that changes in central carbon metabolism impact Wnt signaling [3–6] and the period of the segmentation clock [7], which controls the timing of axis segmentation. Here, we reveal that glycolysis tunes the segmentation clock period in an anti-correlated manner: higher glycolytic flux slows down the clock, and vice versa. Transcriptome and gene regulatory network analyses identified Wnt signaling and specifically the transcription factor Tcf7l2, previously associated with increased risk for diabetes [8, 9], as potential mechanisms underlying flux-dependent control of the clock period. Critically, we show that deletion of the Wnt antagonist Dkk1 rescued the slow segmentation clock phenotype caused by increased glycolysis, demonstrating that glycolysis instructs Wnt signaling to control the clock period. In addition, we demonstrate metabolic entrainment of the segmentation clock: periodic changes in the levels of glucose or glycolytic sentinel metabolite fructose 1,6-bisphosphate (FBP) synchronize signaling oscillations. Notably, periodic FBP pulses first entrained Wnt signaling oscillations and subsequently Notch signaling oscillations. We hence conclude that metabolic entrainment has an immediate, specific effect on Wnt signaling. Combined, our work identifies a glycolysis-FBP-Wnt signaling axis that tunes developmental timing, highlighting the instructive signaling role of metabolism in embryonic development.

## 1 Introduction

Central carbon metabolism impacts gene expression and signal transduction via modulating epigenetic and protein post-translational modifications, while exerting its bioenergetic function by producing energy, reducing equivalents, and cellular building blocks to fuel biological processes [1, 2, 10–14]. While such a widespread role of metabolism is well-known, how metabolism acts as an instructive rather than a permissive signal to control phenotypic outcomes remains a central question. In the definition we use, an instructive signal is information-rich, hence having the capability of tuning a phenotypic outcome, as opposed to a permissive signal leading to a binary effect [15, 16].

To reveal tunability, it is crucial to be able to tune metabolism dynamically and to monitor its impact, for instance at the level of signaling,in real time and in a quantitative manner. Such an approach is applicable to the study of vertebrate embryo mesoderm segmentation. Presomitic mesoderm (PSM) is segmented into somites, the precursors for vertebrae and skeletal muscles, in a periodic fashion [17]. The timing of this process is tightly regulated by a molecular oscillator known as the segmentation clock, which is best characterised by oscillatory activity of the Notch signaling pathway [18]. Temporal periodicity of Notch signaling oscillations is translated into spatial periodicity of somites by integrating additional information encoded by graded signaling pathways such as Wnt, FGF, and retinoic acid [17, 19–21]. In the mouse PSM, FGF and Wnt signaling pathways are also the components of the segmentation clock, exhibiting oscillatory activities coupled to Notch signaling oscillations[19, 22, 23]. Importantly, this highly complex network of interconnected signaling pathways can be dynamically perturbed and functionally studied by using a combination of quantitative liveimaging and a dynamical systems approach. For instance, using microfluidics-based entrainment,we previously showed that the segmentation clock network can be efficiently controlled via external periodic pulses of Notch and Wnt signaling cues,achieving synchronization and tuning of signaling oscillation period [19, 24].

Here, we build on the quantitative live imaging, genetics, and entrainment approach that provide a powerful experimental framework to tackle the central question of how metabolism plays an instructive role. In the PSM, changes in central carbon metabolism impact Wnt signaling [3–6] and the period of the segmentation clock [7]. In particular, glycolysis has been shown to establish an activity gradient from the posterior to anterior PSM [3, 4], being functionally linked to graded signaling activity within the mouse PSM [4–6]. Furthermore, it has been shown that active glycolysis is required for maintaining the segmentation clock oscillation [3]. In this work we addressed whether and how glycolysis plays an instructive role in regulation of developmental timing of mammalian embryo segmentation.

## 2 Results

### 2.1 SSHHGlycolytic flux tunes the period of the segmentation clock

We first asked whether changes in glycolytic flux would have any effect on the segmentation clock period. To manipulate glycolytic flux using genetics, we utilised a conditional cytoPFKFB3 (here-after termed as TG) transgenic mouse line that we previously generated [6]. In this TG line, a cytoplasmic, dominant active form of the glycolytic enzyme PFKFB3 [25] is expressed from the Rosa26 locus upon CRE-recombination, leading to a glucose-dose dependent increase of glycolytic flux in PSM explants [6]. To quantify the segmentation clock period using real-time imaging, we used a fluorescent reporter mouse line, which reflects the oscillatory gene activity of Notch-target gene *Lfng* [26].

Using this experimental strategy, we found that in TG explants cultured in 2.0 mM glucose, the segmentation clock slowed down by about 20% compared to control explants, without arrest of segmentation clock or morphological segmentation defects (Fig. 1A, 1B, Supplementary Video 1). The slowing down of segmentation clock oscillations was also evident when using a Wnt reporter line, i.e., *Axin2-Achilles* knock-in reporter [24] (Extended Data Fig. 1, Supplementary Video 2). To test whether the observed effect on the segmentation clock oscillations is indeed due to an increased glycolytic flux, and not merely the effect of the overexpression of cytoPFKFB3 protein *per se*, we cultured TG explants in reduced glucose concentrations, in order to reduce glycolytic flux (Extended Data Fig. 2A). Indeed, lowering glucose concentration rescued the clock period phenotype in TG explants (Fig. 1B), indicating that the segmentation clock period responds to glycolytic flux rather than cytoPFKFB3 protein *per se*.

**Fig. 1.**
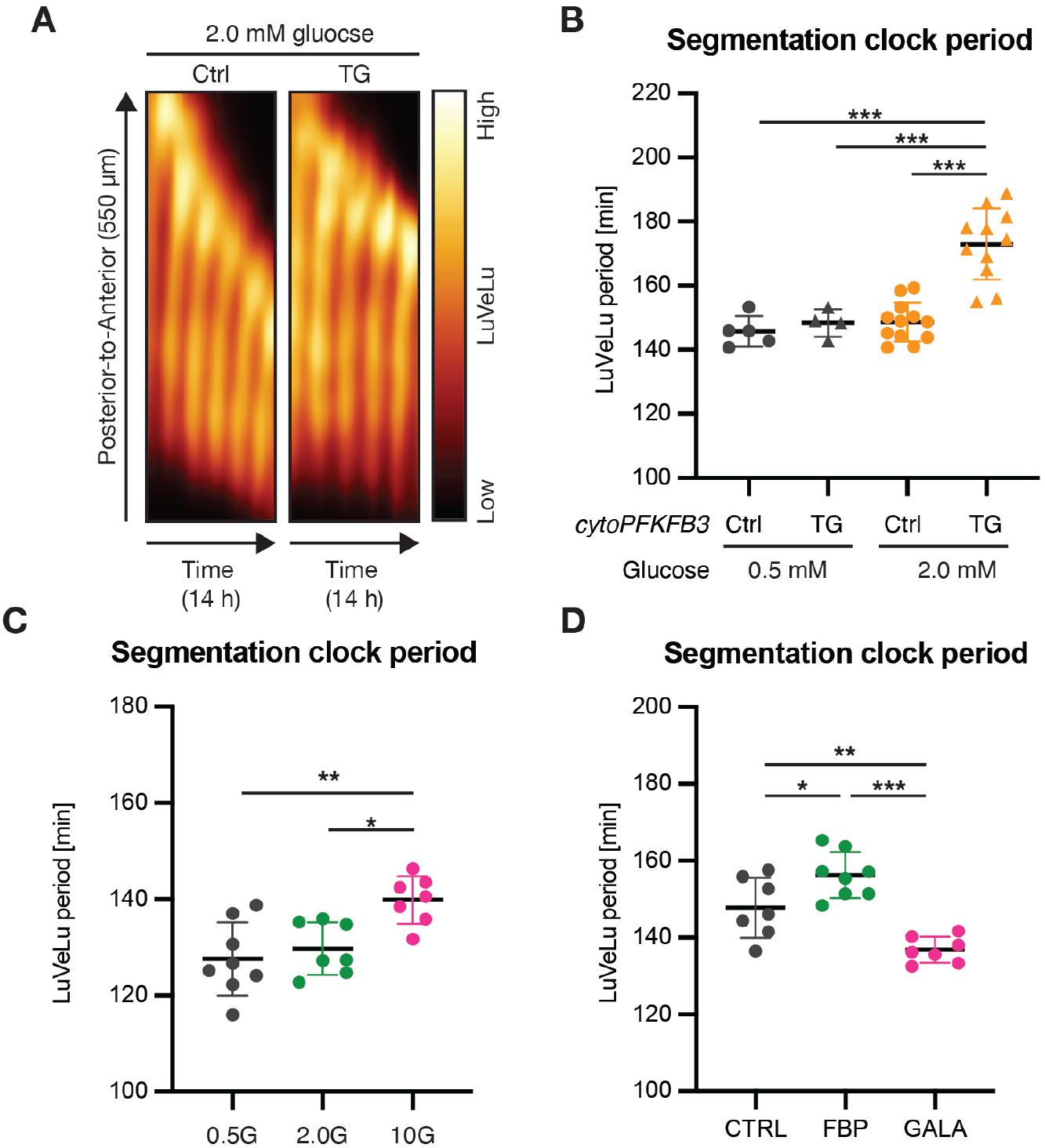
Glycolytic flux tunes the segmentation clock period in an anti-correlated manner. **(A)** Kymographs showing the dynamics of the Notch signaling reporter (= LuVeLu [26]) in control (Ctrl) and cytoPFKFB3 (TG) PSM explants in 2.0 mM glucose condition. **(B-D)** Quantification of the segmentation clock period in various metabolic conditions. The clock periods were determined as a mean of LuVeLu periods between 400-600 min of the imaging. Since the clock period is highly sensitive to temperature, the comparisons are always made within each experiment. (B) The clock period in TG and Ctrl explants cultured in 0.5 mM or 2.0 mM glucose. (C) Effects of glucose titration on the clock period in wild-type explants [0.5 mM (0.5G) vs. 2.0 mM (2.0G) vs. 10 mM (10G) glucose]. (D) Effects of fructose 1,6-bisphosphate (FBP) or galactose (GALA) on the clock period in wild-type explants [CTRL, culture medium with 2.0 mM glucose; FBP, culture medium with 2.0 mM glucose and 10 mM FBP; GALA, culture medium with 2.0 mM galactose (without glucose)]. One-way ANOVA with Tukey’s post hoc test (*p *<*0.05, **p *<*0.01, ***p *<*0.001). Mean ± standard deviation (SD) are shown in the graph, and individual data points represent biological replicates.

To further probe whether glycolytic flux instructs the segmentation clock period, we investigated the impact of tuning (i.e., increasing and decreasing) glycolytic flux in wild-type explants. Importantly, we found that the segmentation clock period was tunable in wild-type explants by modulating glycolytic flux. Increasing glucose led to a slower segmentation clock (Fig. 1C), which we also observed when fructose 1,6-bisphosphate (FBP), a glycolytic sentinel metabolite [6, 27], was supplemented to the medium (Fig. 1D). On the other hand, replacing glucose with galactose, which leads to minimum glycolytic flux (Extended Data Fig. 2B) [28], resulted in the acceleration of the segmentation clock (Fig. 1D). Therefore, minimizing glycolytic flux speeds up the segmentation clock, while increasing glycolytic flux has the opposite effect.

Taken together, our data shows that glycolytic flux tunes the segmentation clock period in an anti-correlated manner.

### 2.2 Characterizing glycolytic flux-induced transcriptional responses in PSM cells

To gain insight into the mechanism underlying the glycolytic flux-dependent control of the segmentation clock period, we next looked into flux-induced transcriptional responses and their potential mechanisms operating in the PSM.

First, we built PSM-specific enhancer-mediated gene-regulatory network (eGRN) using the GRaNIE (Gene Regulatory Network Inference including Enhancers) method [29], which constructs eGRN based on co-variation of chromatin [i.e., transcription factor (TF) binding site] accessibility, TF expression and corresponding target gene expression across samples. We generated paired transcriptome [i.e., RNA sequencing (RNA-seq)] and chromatin accessibility [i.e., assay for transposase-accessible chromatin with sequence (ATAC-seq)] data from wild-type, non-cultured PSM tissues. The PSM tissues were microdissected into tailbud, posterior PSM, anterior PSM, and somite regions, so that a resulting eGRN is linked to gene expression changes following PSM cell differentiation along the embryonic axis, which also mirrors metabolic state changes [3, 4] (Extended Data Fig. 3A).

The resulting eGRN includes 2522 genes out of 28629 (= 9%) genes expressed in the PSM and consists of 69 regulons, where each regulon represents a set of target genes regulated by a TF through their accessible enhancer regions (Extended Data Fig. 3A). These regulons include those associated with TFs that regulate PSM cell differentiation, such as Cdx2 [30] and T [31], providing evidence for the validity of the PSM-specific eGRN inferred with the GRaNIE method.

For the identification of glycolytic flux-responsive genes, we performed transcriptome analysis using explants from control and TG explants cultured in different glucose concentrations for three hours. We limited our analysis to the tailbud region, where the clock period phenotype is most apparent. Combined with the dataset from our previous study [6], this analysis revealed 617 flux-responsive differentially expressed genes (DEGs) that were either upregulated (Cluster (C) 1 and C3) or downregulated (C2, C4, and C5) by increasing glycolytic flux (Fig. 2A, Supplementary Table 1).

**Fig. 2.**
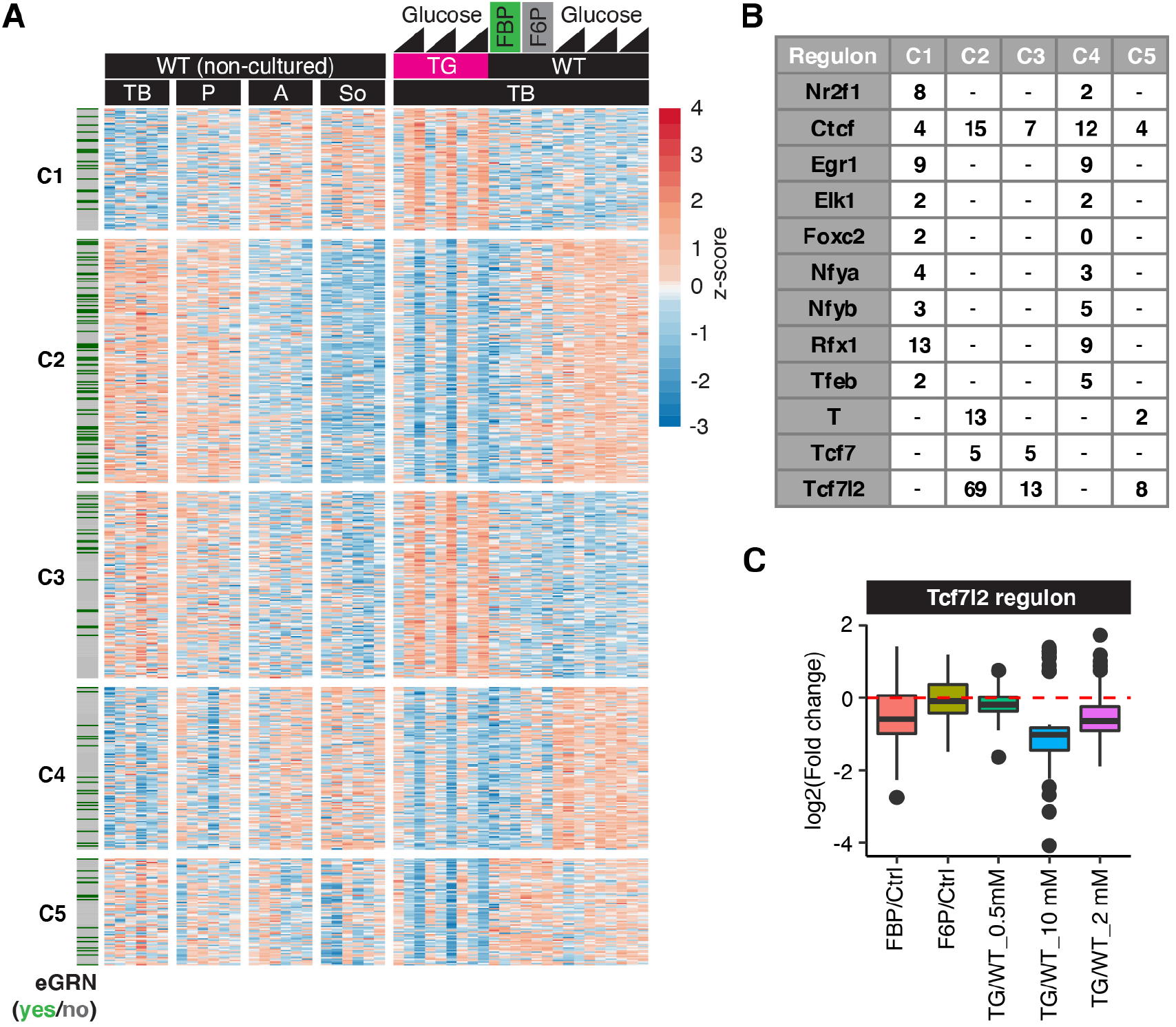
Tcf7l2 regulon responds to glycolytic flux changes within PSM cells. **(A)** A heatmap showing glycolytic flux-responsive differentially expressed genes (DEGs) between wild-type (WT) and cytoPFKFB3 (TG) PSM explants cultured for three-hour in various (i.e., 0.5 mM, 2.0 mM, and 10 mM) glucose conditions (adjusted *p*-value *<*0.01, WT vs. TG for each glucose condition). Normalized counts by variance stabilizing transformation (VST) were used to calculate the z-scores. The datasets were integrated with the datasets from Miyazawa *et al*. (2022) [6]. DEGs that are parts of the PSM-specific eGRN are marked by green. **(B)** A table showing the number of the flux-responsive DEGs that are included in each PSM-specific regulon. **(C)** A box plot showing fold changes in gene expression of the flux-responsive Tcf7l2 targets between different metabolic conditions.

By matching the flux-responsive DEGs to the PSM-specific eGRN, we revealed that 132 DEGs are part of the regulons. Intriguingly, the vast majority of those (90 out of 132 DEGs) are part of the Tcf7l2 regulon (Fig. 2B, Extended Data Fig. 3B). Gene expression of the Tcf7l2 regulon is downregulated with both increased glycolytic flux and FBP supplementation (Fig. 2C, Extended Data Fig. 3C), conditions that we found to cause slowing down of the segmentation clock (Fig. 1).

Tcf7l2 is tightly linked to Wnt signaling [32, 33], and identified as a repressor in our eGRN analysis (Extended Data Fig. 3A). Therefore, these results reveal a glycolysis-Wnt-signaling axis where increased glycolytic flux activates the Tcf7l2 regulon, providing the mechanistic basis for the anti-correlation between glycolytic flux and Wnt signaling target gene expression. Functionally, the glycolysis-Wnt-signaling axis could hence underlie the observed tuning of segmentation clock period.

### 2.3 Glycolysis-Wnt signaling axis controls the segmentation clock period

To functionally test whether the glycolysis-Wnt-signaling axis underlies the flux-dependent tuning of the segmentation clock period, we performed a genetic rescue experiment using a mutant for *Dickkopf-1* (*Dkk1*) [34, 35], a developmentally critical Wnt signaling inhibitor that acts at the level of ligand-receptor interaction. We asked whether partial deletion of *Dkk1* could rescue the clock period phenotype observed in TG embryos, where elevated glycolytic flux correlated with Wnt signaling downregulation. Excitingly, we indeed found that in TG embryos in which one allele of *Dkk1* was deleted, the segmentation clock period was rescued in most of the samples (Fig. 3A). Critically, we found that lactate secretion was not affected by *Dkk1* heterozygosity (Fig. 3B). TG explants maintained high glycolytic flux even in a *Dkk1* heterozygous background, despite showing a rescued clock phenotype. These findings indicate that the proximate cause of the observed clock phenotype in TG embryos are changes in signaling, rather than cellular metabolic state.

**Fig. 3.**
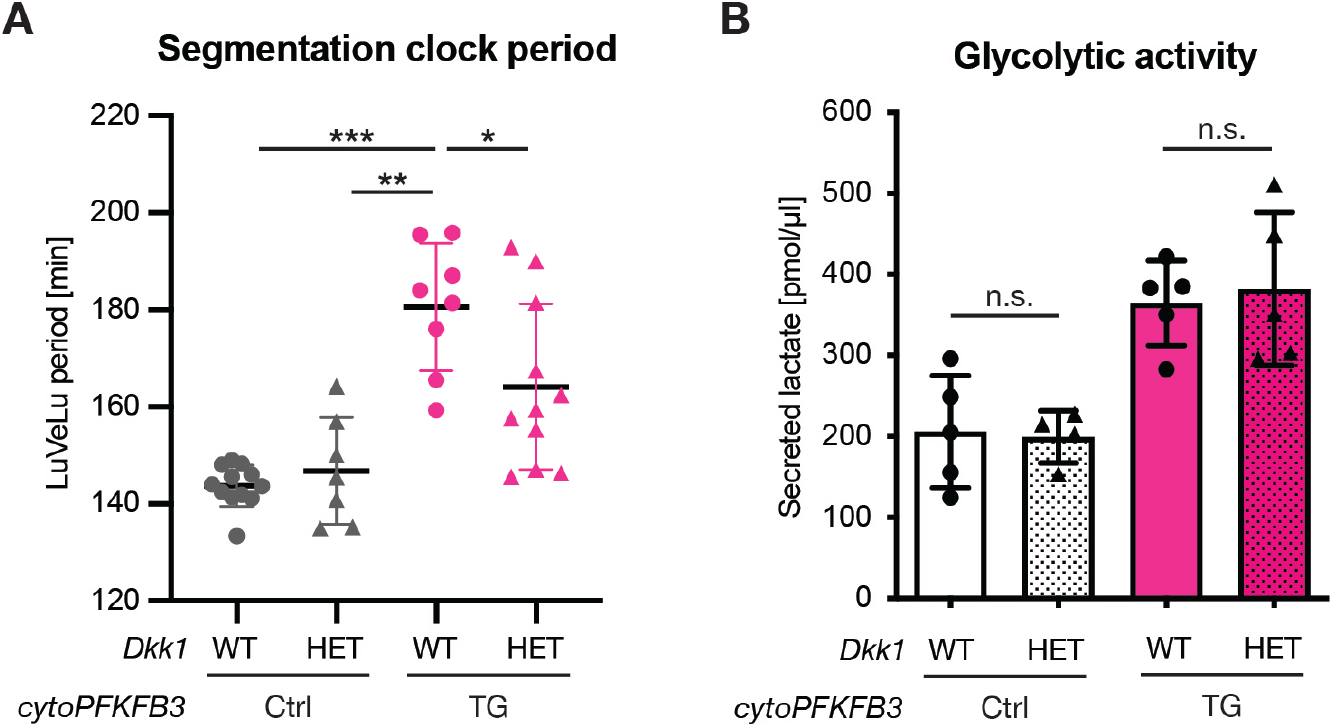
Genetic rescue of the slow segmentation clock phenotype in cytoPFKFB3 embryos without affecting glycolytic flux. **(A)** Quantification of the segmentation clock period in control (Ctrl) and cytoPFKFB3 (TG) explants with one allele of *Dkk1* (HET), compared to samples with wild-type *Dkk1* copy number (WT). The clock period under 2.0 mM glucose condition was determined as a mean of LuVeLu periods between 400-600 min of imaging. One-way ANOVA with Tukey’s post hoc test (*p *<*0.05, **p *<*0.01, ***p *<*0.001). Mean ± SD are shown in the graph, and individual data points represent biological replicates. **(B)** Lactate secretion was quantified as a proxy for glycolytic flux within PSM cells. After 12 h *ex vivo* culture in 2.0 mM glucose, the amount of lactate secreted from control (Ctrl) and cytoPFKFB3 (TG) PSM explants was quantified in wild type (WT) samples with normal *Dkk1* copy number and in samples with one allele of *Dkk1* (HET). Welch’s unpaired t-test (n.s., not significant). Mean ± SD are shown in the graph, and individual data points represent biological replicates.

To further probe the mechanism underlying the clock period phenotype, we also examined whether there is a correlation between cellular redox state and the segmentation clock period, as recently suggested in an embryonic stem cell (ESC)-based model for the segmentation clock [7].

To this end, we quantified the NAD^+^/NADH ratio in control and TG explants under different culture conditions. As expected, the NAD^+^/NADH ratio changed in response to alterations in glycolytic flux (Extended Data Fig. 4). Importantly however, the NAD^+^/NADH ratio was comparable between control explants cultured in 10 mM glucose and TG explants cultured in 2.0 mM glucose (Extended Data Fig. 4), which showed a significant difference in the segmentation clock period (Fig. 1).

Taken together, these data provide strong evidence that the tuning effect of glycolytic flux on the segmentation clock period is not mediated via changes in cellular bioenergetic state but rather, via modulation of Wnt signaling.

### 2.4 Metabolic entrainment of the segmentation clock

To further investigate how glycolytic flux is linked to oscillatory signaling and the segmentation clock, we used a dynamical systems approach based on entrainment. Entrainment offers a quantitative and non-disruptive approach to reveal functional dependencies within a dynamical system. We had previously established microfluidics-based entrainment of the mouse embryo segmentation clock, using periodic pulses of signaling pathway modulators, such as a Notch signaling inhibitor and a Wnt signaling activator [19, 24]. Based on our finding of a functional glycolysis-Wnt-signaling axis, we wondered whether the segmentation clock network could also be entrained by periodic changes in glycolytic flux.

As glycolytic flux in PSM cells can be controlled via the concentration of glucose in the culture medium (Extended Data Fig. 2), we used microfluidics to implement periodic changes in glucose concentration during the culture of PSM explants and monitored segmentation clock dynamics using real-time imaging of a Notch signaling reporter. Strikingly, we found that periodic alternations of glycolytic flux are indeed sufficient to entrain Notch signaling oscillations underlying the segmentation clock (Fig. 4A, 4A’, 4B, 4B’, Supplementary Video 3). We quantified entrainment based on phase-locking (Fig. 4A’, 4B’, Extended Data Fig. 5B) and also using the first Kuramoto order parameter (Fig. 4A, 4B), which effectively measures how synchronous different samples are oscillating.

**Fig. 4.**
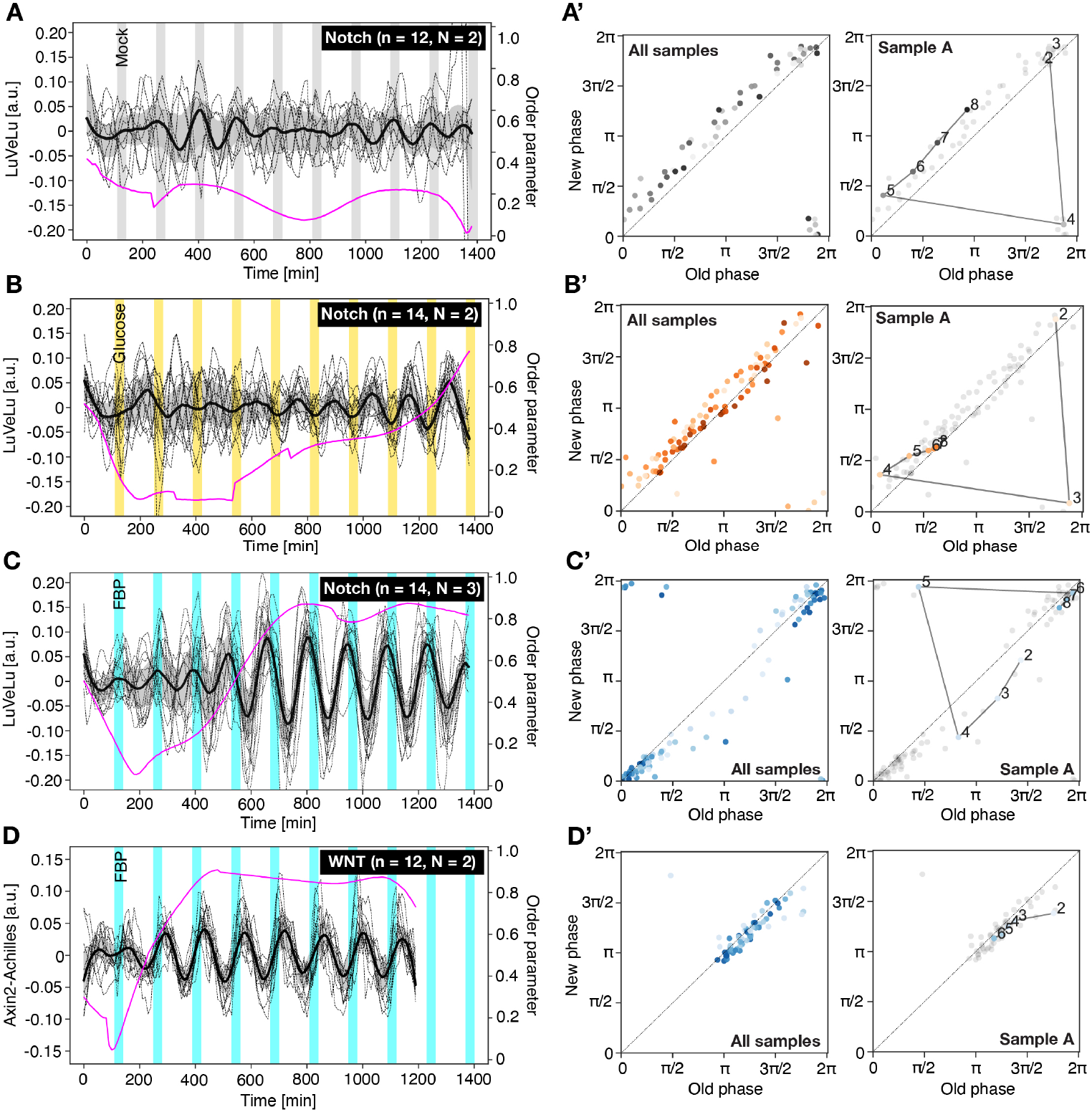
Metabolic entrainment of the segmentation clock. **(A-D)** Detrended (via sinc-filter detrending, cut-off period = 240 min) time-series of LuVeLu (A-C) and Axin2-Achilles (D) intensity oscillations in wild-type PSM explants during metabolic entrainment (dashed lines: individual samples, bold black line: median values, grey shades: the first to third quartile range). Changes in the first Kuramoto order parameter are shown in magenta. Samples were incubated either in a constant (i.e., 2.0 mM) glucose condition with periodic mock pulses (gray) (A) or alternating culture conditions (B-D) with a period of 140-min and a pulse length of 30-min [alternating between: (B) 2.0 mM (white) to 0.5 mM (yellow) glucose conditions; (C, D) the medium with (cyan) or without (white) 20 mM FBP on top of 2.0 mM glucose]. To keep molarity of the medium at constant during experiments, non-metabolizable glucose (i.e., 3-O-methyl-glucose) was added to the medium when necessary. **(A’-D’)** Stroboscopic maps showing step-wise changes in the phase of LuVeLu (A’-C’) and Axin2-Achilles oscillations (D’) during metabolic entrainment. At each pulse of metabolic perturbations with glucose (A’, B’) or FBP (C’, D’), the phase of the oscillator (i.e., new phase) is plotted against its phase at the previous pulse (i.e., old phase). Darker dots represent later time points. Stroboscopic maps of a single representative sample are shown on the right (the numbers in the plots indicate the number of the pulses).

In addition to periodic changes in glucose, we also tested whether periodic pulses of the sentinel metabolite FBP would be sufficient to entrain the segmentation clock. Indeed, our results revealed evidence for Notch signaling entrainment by periodic application of FBP (Fig. 4C, 4C’, Extended Data Fig. 5B, Supplementary Video 4). In contrast, periodic application of pyruvate, the end product of glycolysis, was not sufficient to entrain the segmentation clock (Extended Data Fig. 5A, 5A’, 5B, Supplementary Video 5). These results show that transient, periodic perturbations of glycolysis, specifically at the level of the sentinel metabolite FBP, can entrain the segmentation clock. This provides additional, independent support for glycolytic flux-signaling closely linked to developmental timing.

Importantly, we used metabolic entrainment to further disentangle the functional dependencies between glycolysis, Wnt and Notch signaling pathways. Do periodic FBP pulses entrain Wnt signaling directly or indirectly through Notch signaling entrainment? We previously had shown that Wnt and Notch signaling oscillations are coupled within the segmentation clock network [19]. This means that entrainment of Notch signaling oscillations eventually leads to entrainment of Wnt signalling oscillations with a time delay, and *vice versa*. Thus, we next quantified the timing of metabolic entrainment in regard to both Notch and Wnt signaling oscillations, in order to distinguish direct from more indirect dependencies between glycolysis and Wnt signaling. Notably, we found that periodic FBP pulses first entrained Wnt signaling oscillations, while entrainment of Notch signaling oscillations followed with considerable delay (Fig. 4C, 4D, Supplementary Video 6). Hence, this dynamic entrainment analysis provides strong evidence that glycolysis/FBP has a direct effect on Wnt signaling within the segmentation clock network.

Combined, we show for the first time metabolic entrainment of the segmentation clock, which further establishes a signaling function of glycolysis. Moreover, our analysis of entrainment dynamics supports a specific, direct functional connection of glycolytic flux-signaling to the Wnt signaling pathway.

## 3 Discussion

### 3.1 Glycolysis-FBP-Wnt signaling axis within the PSM

In this study, we show that glycolytic flux tunes the timing of axis segmentation through its instructive function on Wnt signaling. This is supported by our finding that in conditions of increased glycolytic flux, the partial deletion of *Dkk1* rescued the segmentation clock period (Fig. 3). Previously, several mechanisms have been proposed regarding how glucose metabolism impacts Wnt signaling via post-translational modifications [5, 36, 37]. Our results presented here reveal a key signaling role for the glycolytic sentinel metabolite FBP. We propose that the ‘glycolysis-FBP-Wnt signaling axis’ is a module that connects metabolism, signaling and developmental timing. More specifically, combined with our previous study [6], we provide *in vivo* evidence that glycolysis controls Wnt signaling in a dose-dependent, anti-correlated manner (Fig. 1-3). Hence, while increasing glycolytic flux leads to a decrease in Wnt-signaling target gene expression and a slowing down of segmentation, we also see evidence for the inverse: decreasing glycolytic flux within a physiological range correlates with increased Wnt target gene expression and accelerated segmentation. Furthermore, we showed that periodic application of FBP first synchronizes Wnt signaling oscillations and subsequently Notch signaling oscillations during metabolic entrainment (Fig. 4). These findings indicate that glycolytic flux, or its dynamics, tunes Wnt signaling activity to control the timing of the segmentation clock.

These results appear to contrast with findings in studies using *in vitro* stem cell models for mesoderm specification, in which glycolytic inhibition led to downregulation, not upregulation, of Wnt signaling [4, 5, 38–40]. One potential reason for this apparent discrepancy could be rooted in the strength of perturbation applied to glycolytic flux. In the studies mentioned above, glycolysis was either strongly impaired pharmacologically or bypassed altogether (i.e., no glucose condition), which caused downregulation of Wnt (and other signaling) activity. This indeed shows that ongoing glycolysis is required, permissively, for signaling. In contrast, we show that tuning glycolytic flux within the physiological range, both lowering and increasing flux, leads to an anti-correlated response at the level of Wnt signaling targets and segmentation clock period. Combined, the available evidence hence suggest the existence of multiple functional dependencies between glycolysis and signaling. First, a permissive glycolytic function for signaling is evident, i.e., some glycolytic activity is *per se* required. In addition, we show here that an instructive, tunable glycolysis-FBP-Wnt signaling axis exists, controlling the period of the segmentation clock *in vivo*. Future mechanistic studies will further resolve both the permissive and instructive glycolytic function in different contexts.

Intriguingly, during metabolic entrainment, we noticed that periodic changes in glycolytic flux and FBP levels induce periodic changes in tissue shape (Supplementary Video 3-6). This suggests a potential additional link between glycolysis, Wntsignaling, and tissue shape changes. Importantly, however, periodic pulse of pyruvate also caused a similar shape change phenotype but did not result in segmentation clock entrainment. While we therefore conclude that tissue shape changes are not sufficient for entrainment, their link to metabolic signaling needs to be a focus of future studies.

To reveal the detailed mechanisms of glycolytic flux signaling, it will be crucial to identify FBP sensor molecules that mechanistically link intra-cellular FBP levels and Wnt signaling. Probing FBP-protein interactions is one exciting direction that in different contexts have already indicated the widespread regulatory potential of FBP [41– 44]. In addition, our transcriptome and eGRN analyses identified genes within the Tcf7l2 regulon as particularly glycolytic flux-sensitive (Fig. 2). This raises the possibility that Tcf7l2 is a part of the FBP sensing mechanisms and hence FBP-Wnt signaling axis. Notably, Tcf7l2 has been strongly associated with type 2 diabetes and is involved in gluocse homeostatis and insulin secretion in pancreatic *β*-cells [8, 9]. An exciting possibility that requires further investigation is that FBP directly impacts Tcf7l2 activity in an allosteric manner within the segmentation clock network but potentially also in other biological contexts including pancreatic *β*-cells.

### 3.2 Glycolytic flux control of the segmentation clock period

The primary function of the glycolysis-FBP-Wnt signaling axis that we revealed in this study is the control of segmentation clock period in mouse embryos. Previously, Wnt signaling had been functionally linked to the regulation of the segmentation clock period [45], although the underlying mechanisms were not addressed. Our work reveals the direct impact of metabolic state on Wnt-signaling and clock period and hence emphasises the need for future studies to identify how Wnt signaling impacts the period of segmentation clock oscillations. Recently, a series of studies have reported on potential mechanisms of how, in general, the oscillation period can be tuned. Accordingly, a study using *in vitro* stem cell system reported that the segmentation clock period is controlled by mitochondrial respiration, cellular redox state, and ultimately protein translation rate [7]. Additionally, several *in vitro* studies emphasized that differences in protein turnover rates underlie species-specific developmental timing [7, 46–49]. How our *in vivo* findings on the link of glycolysis, Wnt signaling and developmental timing relate to these *in vitro* studies is not resolved yet. In principle, our finding that *increased* glycolytic flux leads to a *slowing* down of the segmentation clock is compatible with a role of mitochondrial respiration, since glycolysis and respiration are considered to be inversely correlated (i.e., Crabtree effect). However, we found that glycolytic flux-signaling shows specificity at the level of FBP, as periodic pulses of pyruvate are not sufficient to entrain the segmentation clock, which could argue against an involvement of mitochondrial respiration. In addition, our findings revealed that glycolysis functions via Wnt signaling (Fig. 3), and not via cellular redox state (Extended Data Fig. 4). We also found clear evidence for a direct immediate effect of FBP on Wnt signaling using metabolic entrainment (Fig. 4). Combined, our findings hence argue against a widespread, bioenergetic mechanism. Instead we identifies a non-bioenergetic metabolic signaling role and reveals the glycolysis-FBP-Wnt signaling axis as a regulator of the segmentation clock period.

### 3.3 Future direction

In conclusion, our study provides evidence that glycolysis is instructive in regulation of Wnt signaling. This regulatory function is crucial for controlling developmental timing and potentially embryonic patterning. The association between glycolysis and Wnt signaling in many biological contexts, ranging from development to disease states [9, 50–52], underscores the critical need to now explore how ubiquitously the glycolysis-FBP-Wnt signaling axis functions in living systems.

These findings also raise the more general question about the significance of the functional link between metabolic activity and developmental timing. We discuss here two, potentially interconnected, hypotheses regarding the broader implications of this relationship.

One appealing hypothesis is that the intrinsic temporal organization of metabolism, which manifests as metabolic rhythms and cycles at various temporal and spatial scales [53], serves as the core template for biological timing and oscillations [54]. In this study, we provide the first demonstration that if present, metabolic cycles (in our case experimentally generated via entrainment) can potently entrain the segmentation clock and developmental timing. Thus, as a next logical step, efforts need to be intensified to elucidate the presence of metabolic rhythms and cycles in living systems.

In addition, the link between metabolism, developmental signaling and timing could serve the integration of environmental cues, such as changes in nutritional resources. Interestingly, we show that access to higher glucose concentrations slows down the pace of embryonic segmentation. In order to understand the significance of this functional dependence between metabolism, signaling and timing, it will be critical to study the dynamic interplay of organisms with their natural environment, considering the entire life cycle.

## 4 Methods

### 4.1 Animal work

All animals were housed in the EMBL animal facility under veterinarians’ supervision and were treated following the guidelines of the European Commission, revised directive 2010/63/EU and AVMA guidelines 2007. All the animal experiments were approved by the EMBL Institutional Animal Care and Use Committee (project code: 21–001 HD AA). The detection of a vaginal plug was designated as embryonic day (E) 0.5, and all experiments were conducted with E10.5 embryos.

### 4.2 Mouse lines

The following mice used in this study were described previously and were genotyped using primers described in these references: *Axin2-Achilles* [24], *Hprt*^*Cre*^ [55], *LuVeLu* [26], Rosa26loxP-stop-loxP-cytoPFKFB3 [6], *Dkk1* mutant [35]. While the *Dkk1* mutant line was maintained on C57BL/6j genetic background, the other mouse lines were maintained on CD1 genetic background. For the genetic rescue experiments, the following primers were used to detect the mutant allele of *Dkk1* [56]: forward, 5’-GCT CTA ATG CTC TAG TGC TCT AGT GAC-3’. Reverse, 5’-GTA GAA TTG ACC TGC AGG GGC CCT CGA-3’.

### 4.3 *Ex vivo* culture of PSM explants

Dissection and *ex vivo* culture of PSM explants were performed as described before [6]. In brief, E10.5 embryos were collected in DMEM/F12 (without glucose, pyruvate, glutamine, and phenol red; Cell Culture Technologies) supplemented with 2.0 mM glucose (Sigma-Aldrich, G8769), 2.0 mM glutamine (Sigma-Aldrich, G7513), 1.0% (w/v) BSA (Cohn fraction V; Equitech-Bio, BAC62), and 10 mM HEPES (Gibco, 15360–106). PSM explants were isolated using a micro scalpel (Feather Safety Razor, No. 715, 02.003.00.715) and were cultured in DMEM/F12 supplemented with 0.5–2.0 mM glucose, 2.0 mM glutamine, and 1.0% (w/v) BSA (Cohn fraction V; Equitech-Bio, BAC62) at 37ºC, under 5% CO_2_, 60% O_2_ condition.

### 4.4 Live imaging of Notch and Wnt signaling reporter lines

To monitor Notch and Wnt signaling activity using real-time imaging, *LuVeLu* [26] and *Axin2-Achilles* knock-in [24] reporter lines were utilized, respectively. Following dissection, PSM explants were washed once with culture medium and were transferred into agar wells (600 nm-width, 3% low Tm agarose, Biozyme, 840101) in 4-well slides (Lab-Tek, #155383). Imaging was performed with a LSM780 laser-scanning microscope (Zeiss), at 37ºC, under 5% CO_2_, 65% O_2_ condition. Samples were excited by a 514 nm-wavelength argon laser through 20×Plan-Apochromat objective (numerical aperture 0.8). Image processing was done using the Fiji software [57]. For extracting period and phase of signaling oscillations, wavelet analysis was performed using pyBOAT [58].

### 4.5 NAD^+^/NADH and lactate measurements

PSM explants without somites were cultured for one hour in DMEM/F12 supplemented with varying amounts of glucose or galactose (Sigma, G0750). The explants were flash frozen by liquid N_2_ following one hour *ex vivo* culture and were stored at -80ºC until use. NAD^+^/NADH measurements were performed according to the manufacturer’s instructions (Promega, G9071). In brief, eight explants were lysed in 90 µl of 0.1N NaOH with 0.5% DTAB and were split into two tubes (40 µl per tube). Samples were then incubated at 60ºC for 15 min with or without adding 20 µl of 0.4N HCl for NAD^+^ and NADH measurements, respectively. After neutralisation either by 0.5M Trizma base solution (for NAD^+^ samples) or Trizma-HCl solution (for NADH samples), the lysates were used for NAD^+^/NADH measurements. Lactate measurements were performed as described before [6].

### 4.6 ATAC- and RNA-sequencing analysis

PSM explants of E10.5 wild-type embryos (CD1 genetic background) were microdissected into tail bud, posterior PSM, anterior PSM, and somite regions by micro scalpel in cold PBS. Each tissue region was transferred into a micro well (ibidi, #80486) and mechanically dissociated to a cell suspension in 4.2 µl cold PBS. Finally, 0.7 µl and 3.3 µl cell suspensions were used for RNA-sequencing (RNA-seq) and ATAC-sequencing (ATAC-seq), respectively. For the comparison between control and *cytoPFKFB3* PSM explants, explants were cultured for three-hour *ex vivo* before collecting tail buds for RNA-seq analysis.

#### ATAC-seq

We followed the Omni-ATAC protocol [59] with some modifications. For transposition reactions, 3.3 µl cell suspensions were mixed with 5.0 µl 2x TD buffer (20 mM Tris-HCl pH 7.6, 10 mM MgCl_2_, 20% dimethyl formamide), 1.0 µl TDE1 (Illumina, #15027865), 0.1 µl 1% digitonin (Promega,#G9441), 0.1 µl 10% Tween-20 (Sigma, #11332465001), 0.1 µl 10% NP-40 (sigma, #11332473001), and 0.4 µl nuclease-free water. After 30 min incubation at 37ºC on a thermomixer set at 600 rpm, the samples were purified by a DNA Clean and Concentrator-5 (Zymo Research, D4014) and DNA concentrations were determined by Qubit Fluorometer (dsDNA High Sensitivity Kit, ThermoFisher, Q32851). The samples were diluted to 20 ng/µl and used as templates for library preparations by PCR. PCR reactions were performed using primers from Nextera XT Index Kit (Illumina, FC-131-1001) and NEBNext High Fidelity 2X PCR Master Mix (NEB, M0541). After purification with Qiagen MinElute PCR Purification Kit (Qiagen, 28004), individual libraries were size selected (100–800 bp) with Ampure XP beads (Beckman Coulter, #A63881). Libraries were quantified using the Qubit Fluorometer (dsDNA High Sensitivity Kit) and average fragment length distribution was determined by the Bioanalyzer (Agilent, High Sensitivity DNA kit, 5067-4626). Prepared libraries were multiplexed in pools of equimolar concentrations and sequenced on the NextSeq 500 (Illumina) platform with 75-bp paired-end readings. After demultiplexing and barcode trimming (Trimmomatic Galaxy Version 0.36.6), sequencing reads were quality checked (FastQC Galaxy Version 0.73) and aligned to Mus Musculus genome (GRCm38) with the Bowtie2 aligner (Galaxy Version 2.3.4.2, options -I 0 -X 2000 –dovetail –sensitive). Multi-mapping and duplicate reads were removed; finally only reads mapping to major chromosomes were kept [60].

#### RNA-seq

We followed the Smart-seq2 protocol [61] with some modifications. In brief, dissociated cells were lysed with three times volume of cell lysis buffer (0.02% Triton-X with RNasin), snap frozen by liquid N_2_, and stored at -80ºC until cDNA synthesis. cDNAs were synthesized using SuperScript IV Reverse Transcriptase (Thermo Fisher Scientific) and amplified by PCR with HiFi Kapa Hot start ReadyMix (Kapa Biosystems, KK2601). After clean-up with SPRI beads, concentrations of cDNA (50-9000 bp) samples were determined by the Bioanalyzer (Agilent, High Sensitivity DNA kit). 250 pg cDNAs were then used for tagmentation-based library preparation. Libraries were quantified using the Qubit Fluorometer (dsDNA High Sensitivity Kit) and average fragment length distribution was determined by the Bioanalyzer (Agilent, High Sensitivity DNA kit, 5067-4626). Prepared libraries were multiplexed in pools of equimolar concentrations and sequenced on the NextSeq 500 (Illumina) with 75-bp paired-end (for the wild-type, non-cultured PSM explants) or single-end (for the comparison between control and *cytoPFKFB3* explants) readings. After demultiplexing and barcode trimming (TrimGalore Galaxy Version 0.4.3.1), sequencing reads were quality checked (FastQC Galaxy Version 0.69) and aligned to Mus Musculus genome (GRCm38) with the with the STAR aligner (version 2.5.2b, default options) [60]. Multi-mapping reads were removed and RNA-seq quality assessed with Picard CollectRnaSeqMetrics (Galaxy version 2.7.1.1)

### 4.7 GRaNIE analysis

Enhancer-mediated gene regulatory network(eGRN) was constructed from the matched RNA-seq and ATAC-seq data (24 samples for each) ofthe PSM explants from E10.5 wild-type embryos using the developer’s version of the now published GRaNIE package (https://bioconductor.org/packages/release/bioc/html/GRaNIE.html)[29]. Raw gene counts from RNA-seq data were produced with a summarizeOverlaps function from the GenomicAlignments R package(https://bioconductor.org/packages/release/bioc/html/GenomicAlignments.html) [62], corrected for different experimental batches using Combat-seq function from the R package sva[63] and log2 normalised. ATAC-seq peak counts were generated using DiffBind R package(https://bioconductor.org/packages/DiffBind/),and peak positions were identified using MACS2software (https://genomebiology.biomedcentral.com/articles/10.1186/gb-2008-9-9-r137) [64]. The details of the GRaNIE approach are described here [29]. Briefly, in the first step the expression of each TF was correlated with accessibility of each of the accessible regions (=ATAC-seq peak)with and without a known binding site of theTF (foreground and background, respectively).Known binding sites were defined using theHOCOMOCO database v.10 [65]. Significantly correlated TF-peak links were identified using empirical FDR of 30% (calculated separately foreach TF) and an absolute correlation Pearson’scoefficient of *>*0.4. In the second step chromatin accessibility at the ATAC-seq peaks was correlated with the expression of all genes less than250kb away from the peak and peak-gene links were retained if they were positively and significantly (P *<*0.05) correlated (our assumption i that accessibility at the regulatory region positively correlates with expression of the linked gene), and if their Pearson’s correlation coefficient was *>*0.4. This resulted in the eGRN consisting of 69 TFs, 5154 TF-peak-gene connections of 2522 unique genes. TF regulons were defined as all TF-gene links of each TF within the network.

### 4.8 Microfluidics-based segmentation clock entrainment

PDMS chips and PTFE tubing (inner diameter: 0.6 mm, APT AWG24T) for microfluidics-based entrainment experiments were prepared as described before [19, 24]. Culture media were prepared on the day of experiments by adding a metabolite of interest [either glucose, FBP, pyruvate (Sigma, P4562), or 3-OMG (Sigma, M4879)] to DMEM/F12 supplemented with 2.0 mM glutamine (Sigma-Aldrich, G7513), 0.01% (w/v) BSA (Cohn fraction V; Equitech-Bio, BAC62), and 1% penicillin-streptomycin (Gibco, 15140122). The PDMS chip (soaked in PBS) and the culture medium (filled in 10 mL syringes; BD Biosciences, 300912, diameter 14.5 mm) were degassed before use for at least one hour in a vacuum desiccator chamber.

Following dissection, PSM explants with two intact somites were transferred to the PDMS chip and sample inlets were plugged with a PDMS-filled PTFE tubing. The tubings connected to the syringes with medium were then connected to the medium inlets and the samples were placed in the incubator (37ºC, 5% CO_2_, 65% O_2_) installed on a LSM780 laser-scanning microscope (Zeiss) for preculture. Pumping was started for both the control and treatment medium at the flow rate of 20 µl/hr. A half hour later, only the control medium was pumped into the chip for another 30 min at the flow rate 60 µl/hr. After the pre-culture, imaging was started under constant or alternating culture conditions.

For data analysis, moving ROIs (30-pixel in diameter) were placed in the posterior PSM to obtain intensity profiles of LuVeLu or Axin2-Achilles reporters over time. To extract the period and phase of LuVeLu and Axin2-Achilles oscillations, the intensity profiles were analysed using a wavelet analysis workflow [58]. Entrainment of Notch and Wnt signaling oscillations was analysed using stroboscopic maps and the first Kuramoto order parameter as described before [24].

### 4.9 Data availability

The ATAC-seq and RNA-seq data generated in this study were deposited in the BioStudies under the accessioin number E-MTAB-13692, E-MTAB-13693, and E-MTAB-13694. For identifying glycolytic flux-responsive genes, the RNA-seq data from our previous study [available in the European Nucleotide Archive (ENA) under the accession number PRJEB55095] were also used [6].

## Acknowledgments

We thank Irene Miguel-Aliaga, Kristina Stapornwongkul, and Vikas Trivedi for their feedback on the manuscript, Vladimir Benes and Laura Villacorta for technical advice and support for RNA-seq analysis and Jonathan Landry for helping RNAseq data analysis. This work is supported by the EMBL Advanced Light Microscopy Facility (ALMF), Genomics Core Facility, and all the member of Laboratory Animal Resource (LAR). H.M. was supported by the EMBL Interdisciplinary Postdoc (EI3POD) programme under H2020 Marie Sklodowska-Curie Actions COFUND (grant number 664726) and the Japan Society for the Promotion of Science (JSPS). E.E. was supported by the Human Frontier Science Program (HFSP) fellowship. This work was supported by the European Molecular Biology Laboratory and received funding from the European Research Council under an ERC consolidator grant agreement n.866537 to A.A. and the German Research Foundation/DFG (project SFB 1324 – project number 331351713) to A.A.

## 6 Author contributions

H.M.: Conceptualization, Methodology, Formal analysis, Investigation, Writing - Original Draft, Visualization, Supervision

J.R.: Conceptualization, Methodology, Formal analysis, Investigation, Writing - Original Draft, Visualization

P.G.L.S.: Methodology, Software, Investigation

E.E.: Methodology, Formal analysis, Investigation D.B.: Software, Formal analysis, Investigation

C.G.: Software, Formal analysis

J.Z.: Supervision, Funding acquisition

A.A.:Conceptualization, Methodology, Writing – Original draft preparation, Supervision, Project administration, Funding acquisition

## Declarations

The authors declare that they have no conflict of interests.

## Appendix A Extended Data

**Extended Data Fig. 1.**
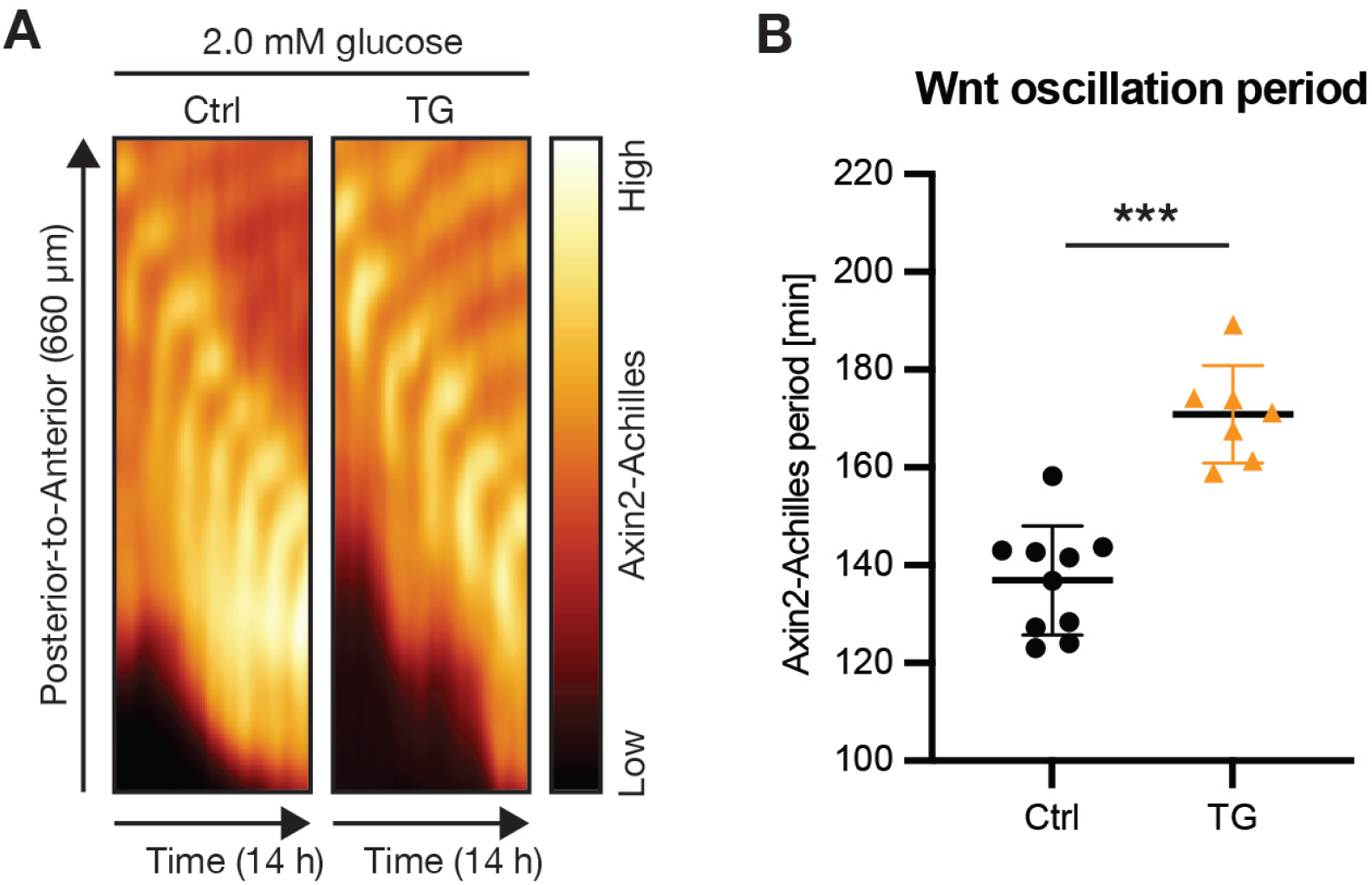
Increasing glycolytic flux slows down Wnt signaling oscillations. **(A)** Kymographs showing the dynamics of the Axin2-Achilles knock-in reporter in control (Ctrl) and cytoPFKFB3 (TG) PSM explants in 2.0 mM glucose condition. **(B)** Quantification of the Wnt signaling oscillation periods in Ctrl and TG explants cultured in 2.0 mM glucose. The periods were determined as a mean of Axin2-Achilles periods between 400-600 min of the imaging. Welch’s unpaired t-test, ***p *<*0.001. Mean ± SD are shown in the graph, and individual data points represent biological replicates.

**Extended Data Fig. 2.**
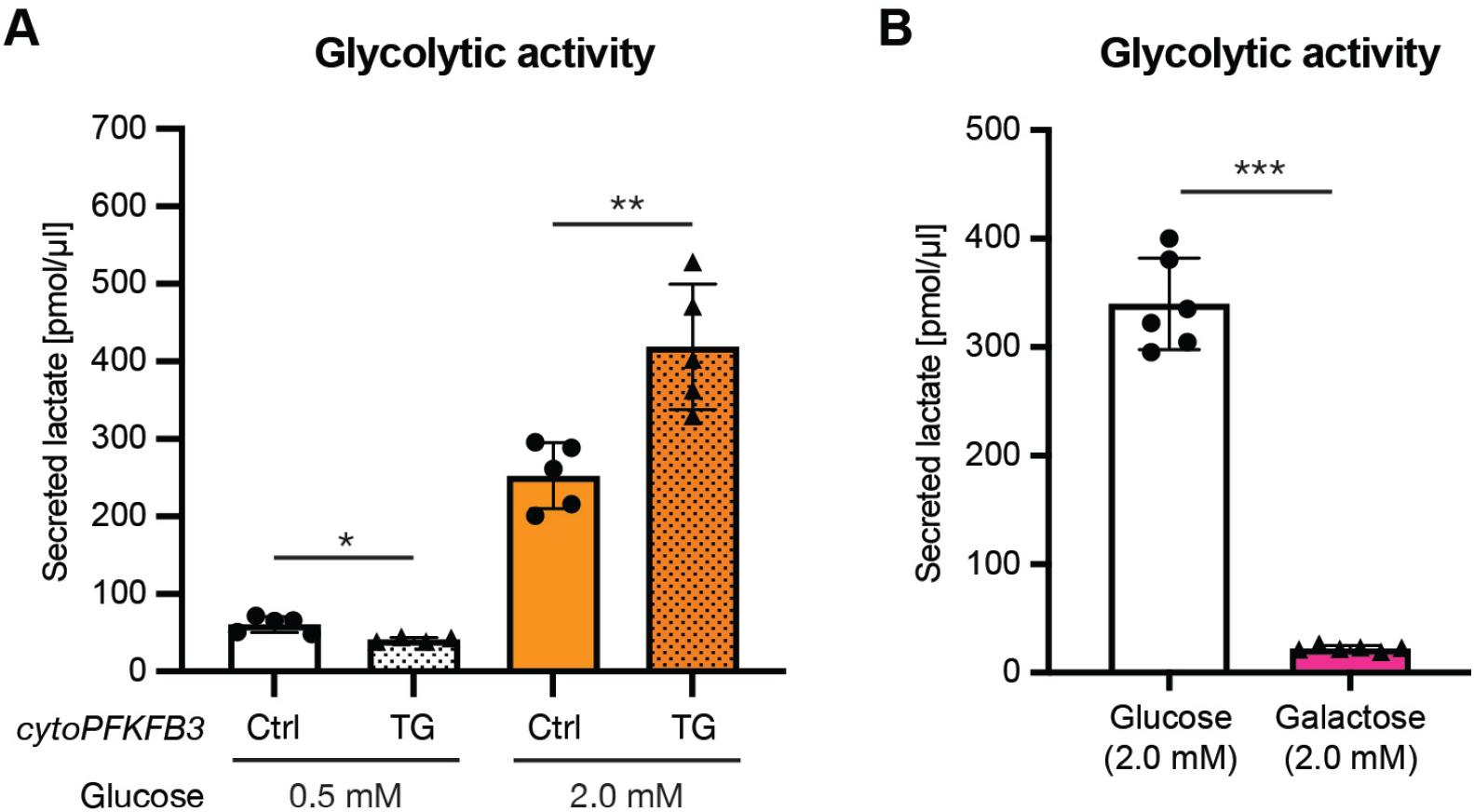
Glycolytic flux shows glucose-dose dependency in PSM cells. **(A, B)** Lactate secretion was quantified as a proxy for glycolytic flux within PSM cells. The amount of lactate secreted from PSM explants during 12 h ex vivo culture was quantified. (A) Comparison of lactate secretion between control (Ctrl) and cytoPFKFB3 (TG) explants cultured in 0.5 mM or 2.0 mM glucose (the data for 2.0 mM glucose condition is adapted from Miyazawa et al. 2022 [6]). (B) The effect of replacing glucose with galactose on lactate secretion from wild-type explants. Welch’s unpaired t-test, *p *<*0.05, **p *<*0.01 vs. Ctrl. Mean ± SD are shown in the graph, and individual data points represent biological replicates.

**Extended Data Fig. 3.**
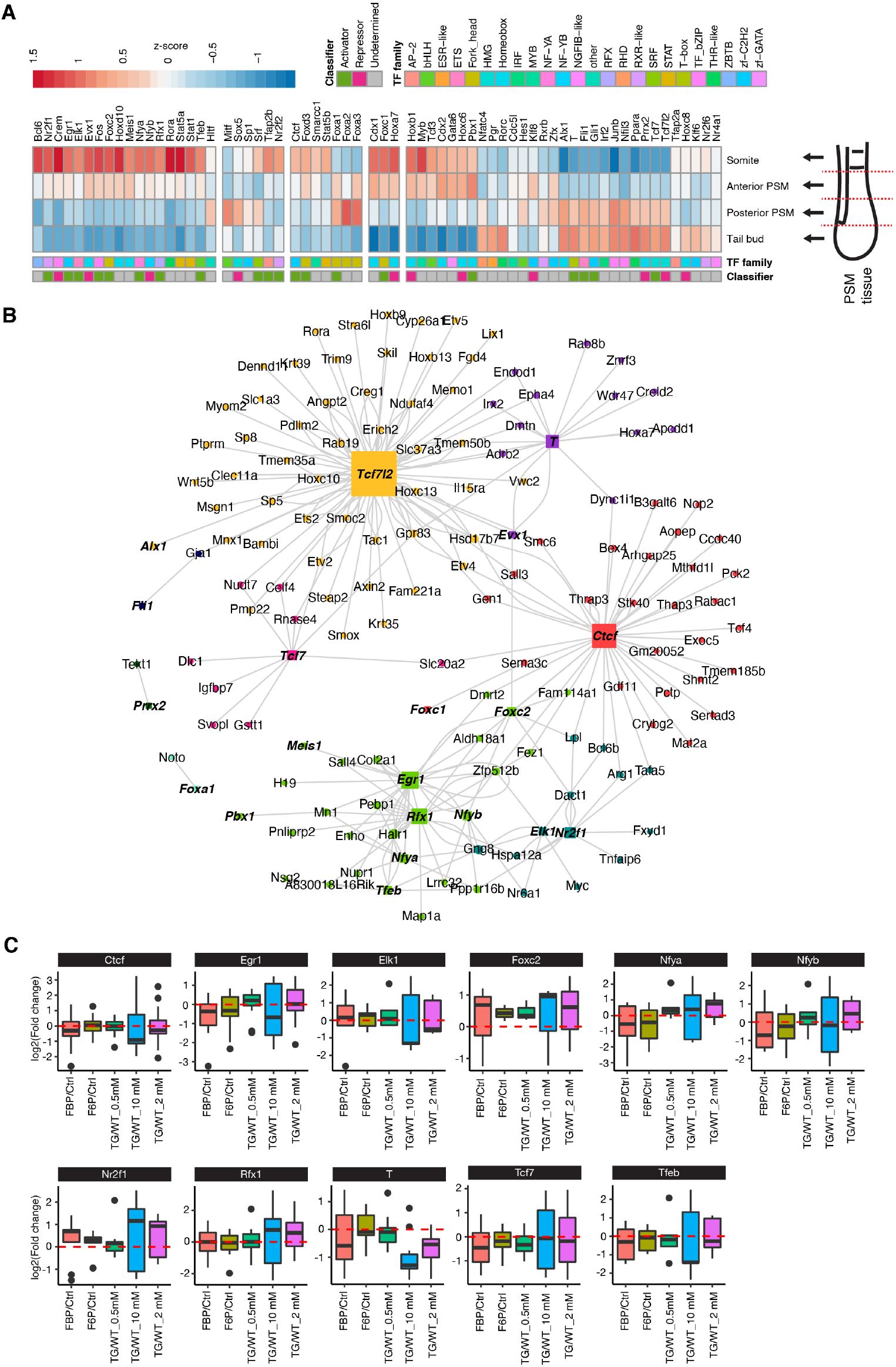
Building a PSM-specific eGRN using the GRaNIE method. **(A)** A heatmap showing gene expressions of each PSM-specific regulon (i.e., means of all the targets) identified by the GRaNIE method. Normalized counts by variance stabilizing transformation (VST) were used to calculate the z-scores. **(B)** A network showing TFs (colored squares) and their glycolytic flux-responsive target genes (colored circles). **(C)** Box plots showing fold changes in gene expressions of flux-sensitive DEGs that constitutes each PSM-specific regulon. The fold changes were calculated between different metabolic conditions.

**Extended Data Fig. 4.**
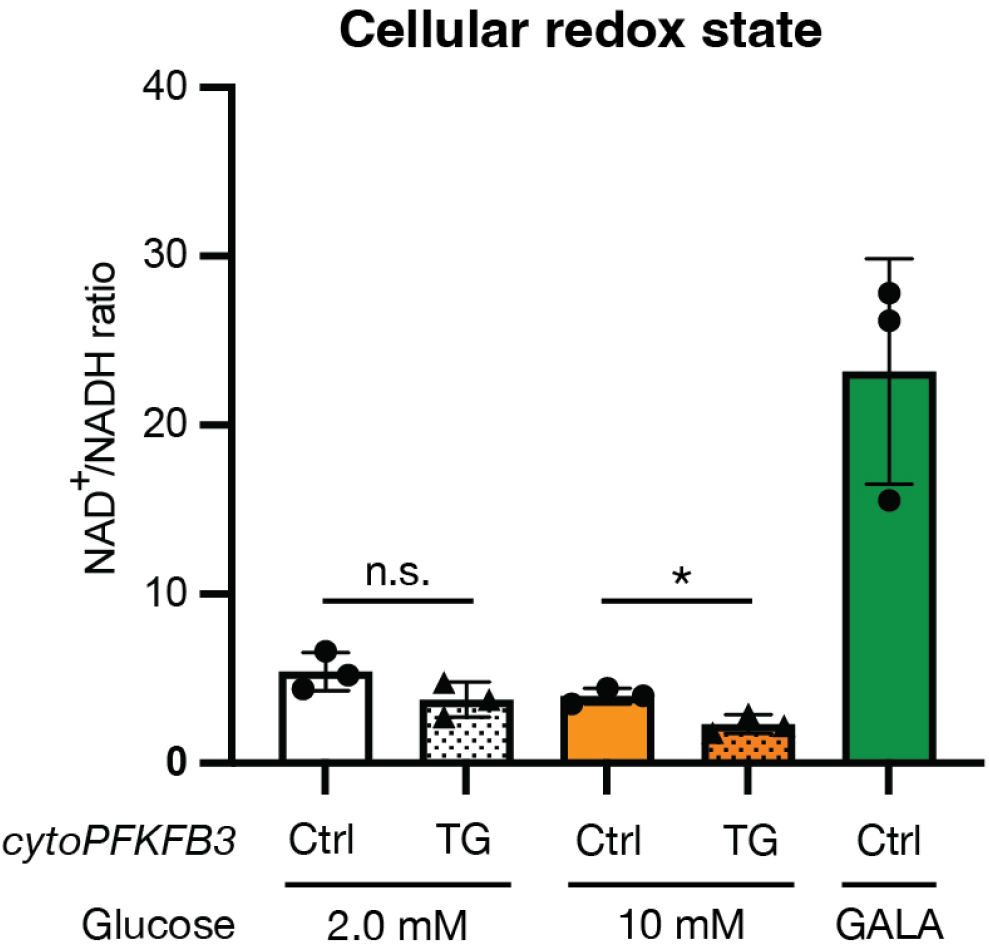
Response of cellular redox state to alterations in glycolytic flux within PSM cells. Quantification of NAD^+^/NADH ratio following one-hour ex vivo culture of control (Ctrl) and cytoPFKFB3 (TG) PSM explants under various culture conditions. For the galactose (GALA) condition, 2.0 mM galactose was supplemented to the culture medium instead of glucose. Mean ± SD are shown in the graph, and individual data points represent biological replicates. Welch’s unpaired t-test, *p *<*0.05.

**Extended Data Fig. 5.**
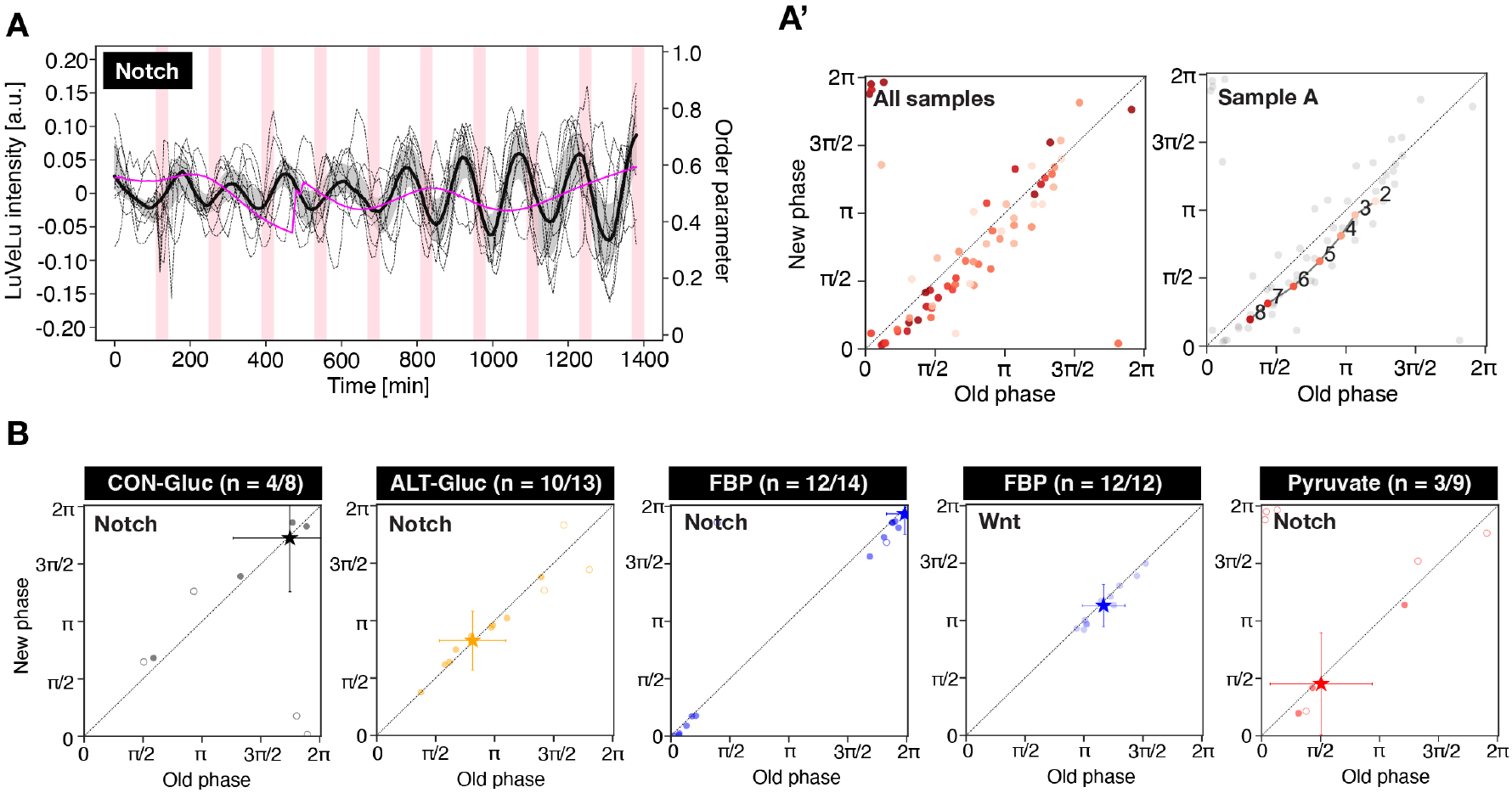
Segmentation clock entrainment by periodic, transient glycolytic cues. **(A)** Detrended (via sinc-filter detrending, cut-off period = 240 min) time-series of LuVeLu intensity oscillations in wild-type PSM explants exposed to periodic pulses of 20 mM pyruvate (dashed lines: individual samples, bold black line: median values, grey shades: the first to third quartile range). Changes in the first Kuramoto order parameter are shown in magenta. To keep molarity of the medium at constant during experiments, 20 mM non-metabolizable glucose (i.e. 3-O-methyl-glucose) was added to the basal medium containing 2.0 mM glucose. **(A’)** Stroboscopic maps showing step-wise changes in the phase of LuVeLu oscillations in response to periodic pyruvate pulses. Darker dots represent later time points (the numbers in the plots indicate the number of the pulses). **(B)** Stroboscopic maps showing the phase of Notch (i.e., LuVeLu) and Wnt (i.e., Axin2-Achilles) oscillations at the last pulse of metabolite. Filled circles represent entrained samples, while open circles represent non-entrained samples. Samples are considered to be entrained when a phase difference between the last and second last pulses is less then *π*/8. CON-Gluc, constant (2.0 mM) glucose condition; ALT-Gluc, alternating (from 2.0 mM to 0.5 mM) glucose condition.

